# Electrochemical DNA-based sensors for measuring cell-generated forces

**DOI:** 10.1101/2023.12.03.569814

**Authors:** Mahmoud Amouzadeh Tabrizi, Priyanka Bhattacharyya, Ru Zheng, Mingxu You

## Abstract

Mechanical forces play an important role in cellular communication and signaling. We developed in this study novel electrochemical DNA-based force sensors for measuring cell-generated adhesion forces. Two types of DNA probes, i.e., tension gauge tether and DNA hairpin, were constructed on the surface of a smartphone-based electrochemical device to detect piconewton-scale cellular forces at tunable levels. Upon experiencing cellular tension, the unfolding of DNA probes induces the separation of redox reporters from the surface of the electrode, which results in detectable electrochemical signals. Using integrin-mediated cell adhesion as an example, our results indicated that these electrochemical sensors can be used for highly sensitive, robust, simple, and portable measurement of cell-generated forces.

## Introduction

Cell-generated adhesion force is a key component in regulating various cellular processes such as the migration, deformation, invasion, and metastasis of cancer cells.^[1]^ During these processes, cell membrane adhesion molecules like integrins and cadherins bind with their target ligands and apply piconewton (pN)-scale forces to mediate cell–extracellular matrix and cell–cell adhesions.^[1,2]^ Several techniques have been developed for the detection of these cellular forces including traction force microscopy,^[3]^ micropillar arrays,^[4]^ atomic force microscopy,^[5]^ optical/magnetic tweezers,^[6]^ and fluorescent tension probes.^[7]^ Although these powerful approaches have significantly improved our ability to measure cell-generated adhesion forces,^[8]^ the requirement of large/expensive instruments and skilled operators are currently limiting their potential adoption by the broader scientific community. On the other hand, we realized that electrochemical sensors can be more portable, inexpensive, and easier-to-use, which once incorporated for measuring cellular forces, would potentially start a new chapter in mechanobiology. We aim to test this idea by developing novel electrochemistry-based cellular force sensors in this project.

To engineer electrochemical force sensors, we consider integrating force-sensitive DNA probes as the detection units for cell-generated adhesion forces. DNAs are arguably the most popularly used moieties for designing molecular tension probes, given their modularly tunable force threshold value, high stability, and ease of synthesis and modification.^[9]^ DNA tension probes are often designed based on either an irreversible double-stranded mode (e.g., so-called tension gauge tether, **TGT**, for detecting forces of ∼12–56 pN)^[10]^ or a reversible **hairpin** mode (with force threshold value F_1/2_ in the range of ∼2–20 pN).^[11]^ Herein, we want to study the possibility of developing both TGT and hairpin DNA tension probes that can generate electrochemical signals upon experiencing different levels of cellular adhesion forces.

In our design (Figure 1), the self-assembled DNA tension probes contain (1) a thiol group at one end to tether to the surface of a gold screen-printed electrode (**Au-SPE**), (2) a cyclic arginine-glycine-aspartic-acid-D-phenylalanine-lysine (cRGDfK or in short **RGD**) ligand at the other end or in another strand to recognize cell surface integrins, i.e., widely used transmembrane receptors for cell adhesion, (3) a methylene blue (**MB**) redox reporter that can undergo reversible electron transfer and release two electrons per redox reaction,^[12]^ and (4) a force-sensitive DNA hairpin (F_1/2_, 12 pN) or a double-stranded TGT structure with low (∼12 pN), medium (∼43 pN), or high (∼56 pN) threshold forces. In both hairpin and TGT modes, in the absence of cellular forces, the distance of MB from the surface of conductive Au-SPE is minimal and as a result, exhibiting large electron transfer rate and high intensity of electrochemical signals. Once the DNA hairpin or TGT duplex is unfolded due to integrin-mediated cell adhesion, an increased distance between MB and Au-SPE will immediately result in a decreased electron transfer and electrochemical signals. In such a way, specific cell-generated adhesion forces can be converted into detectable electrochemical signals.

**Figure 1.**
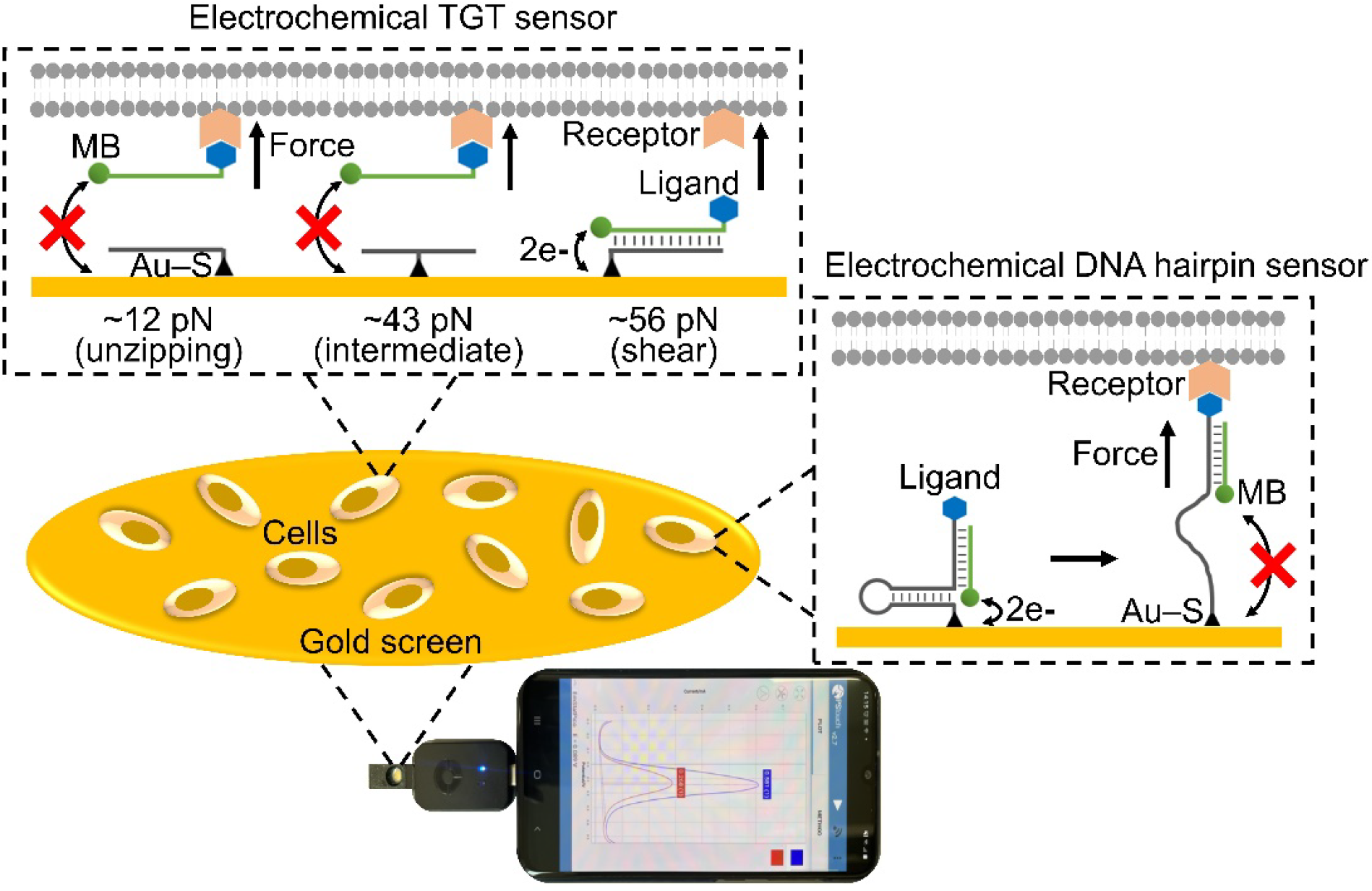
Schematic and photo image of the smartphone-based electrochemical DNA-based force sensors. Thiolated tension gauge tether (TGT) or DNA hairpin probes were immobilized on the surface of a gold screen-printed electrode. Cell adhesion force mediated by a specific receptor– ligand pair could separate the double-stranded TGT probes or unfold the DNA hairpin, and as a result, the attached methylene blue reporter exhibited a decreased electrochemical signal.

By attaching the hairpin- or TGT probe-incorporated Au-SPE chip to a smartphone-based electrochemical device, our results indicate that integrin-mediated forces during the adhesion of HeLa cells can indeed be precisely and sensitively detected by these portable electrochemical DNA-based sensors. To the best of our knowledge, this is for the first time that electrochemical sensors can be developed for the detection of molecular forces during cell adhesions. We expect these new electrochemical DNA-based force sensors can be powerful complementary tools to existing mechanobiology techniques in improving our understanding of cell-generated forces.

## Results

### Construction and characterization of the electrochemical TGT-based force sensors

We first designed three electrochemical TGT force sensors based on previously characterized DNA duplex structures that can be ruptured by forces at ∼12 pN, 43 pN, and 56 pN, respectively.^[10a]^ These TGT probes share the same DNA sequence but have different force application geometries, depending on the relative position of the thiol anchor group on one strand (**anchor strand**) to that of the RGD ligand on the other (**reporter strand**). Using the same 5’-MB-3’-RGD-modified reporter strand (Table S1), by inserting the thiol group at the 5’-end (unzipping mode), middle (intermediate mode), or 3’-end (shear mode) of the anchor strand, low-level, medium-level, and high-level forces can be detected, correspondingly.

To construct these sensors, the thiolated anchor strand was first added and self-immobilized onto surface-cleaned Au-SPE through the formation of Au–S bonds (Figure S1a). After washing away nonspecifically adsorbed anchor DNA and passivating with 1-hexanethiol, 5’-MB-3’-biotin-modified reporter strand was then added and followed by incubation with streptavidin and *biotinylated* cyclic arginine-glycine-aspartic-acid-D-phenylalanine-lysine to finally fabricate TGT probes on the electrode surface.

Electrochemical impedance spectroscopy (EIS)^[13]^ and the corresponding Nyquist plots were used for the stepwise characterization of this fabrication process. In a Nyquist plot, a semicircular portion at high frequencies and a linear portion at low frequencies is related to the charge-transfer-limited process and diffusion-limited process, respectively. By analyzing the diameter of the semicircle, the charge-transfer resistance (**R**_**ct**_) of the electrode can be obtained.^[13]^ Our EIS measurements were performed by applying 0.17 V as a DC voltage, i.e., a formal potential of ferrocene carboxylate, a redox probe used in this test (Figure S1b). Using 12 pN TGT sensor as an example, as can be seen in Figure 2a, the unmodified Au-SPE exhibited a low R_ct_ of ferrocene carboxylate at ∼120 Ω. After the conjugation of the 5’-thiolated anchor strands, the R_ct_ value increased to ∼1850 Ω, which is due to the electrostatic repulsion between the negatively charged ferrocene carboxylate and the phosphate groups of DNA oligonucleotides. We further estimated the surface coverage of the DNA strand (**θ**_**t**_) to be ∼93% by using the equation θ_t_ = (1 – R_ct_/R_ct_′) ×100%,^[14]^ where R_ct_ and R_ct_′ is respectively the charge transfer resistance of bare Au-SPE and that after DNA modification. Upon adding 5’-MB-3’-biotin-modified reporter strand, the R_ct_ of the electrode increased to ∼2510 Ω (Figure 2a), as the formation of double-stranded TGT caused a further limited mass transfer of ferrocene carboxylate to the electrode surface. These EIS-based stepwise characterizations indicated that the TGT sensors were fabricated successfully.

**Figure 2.**
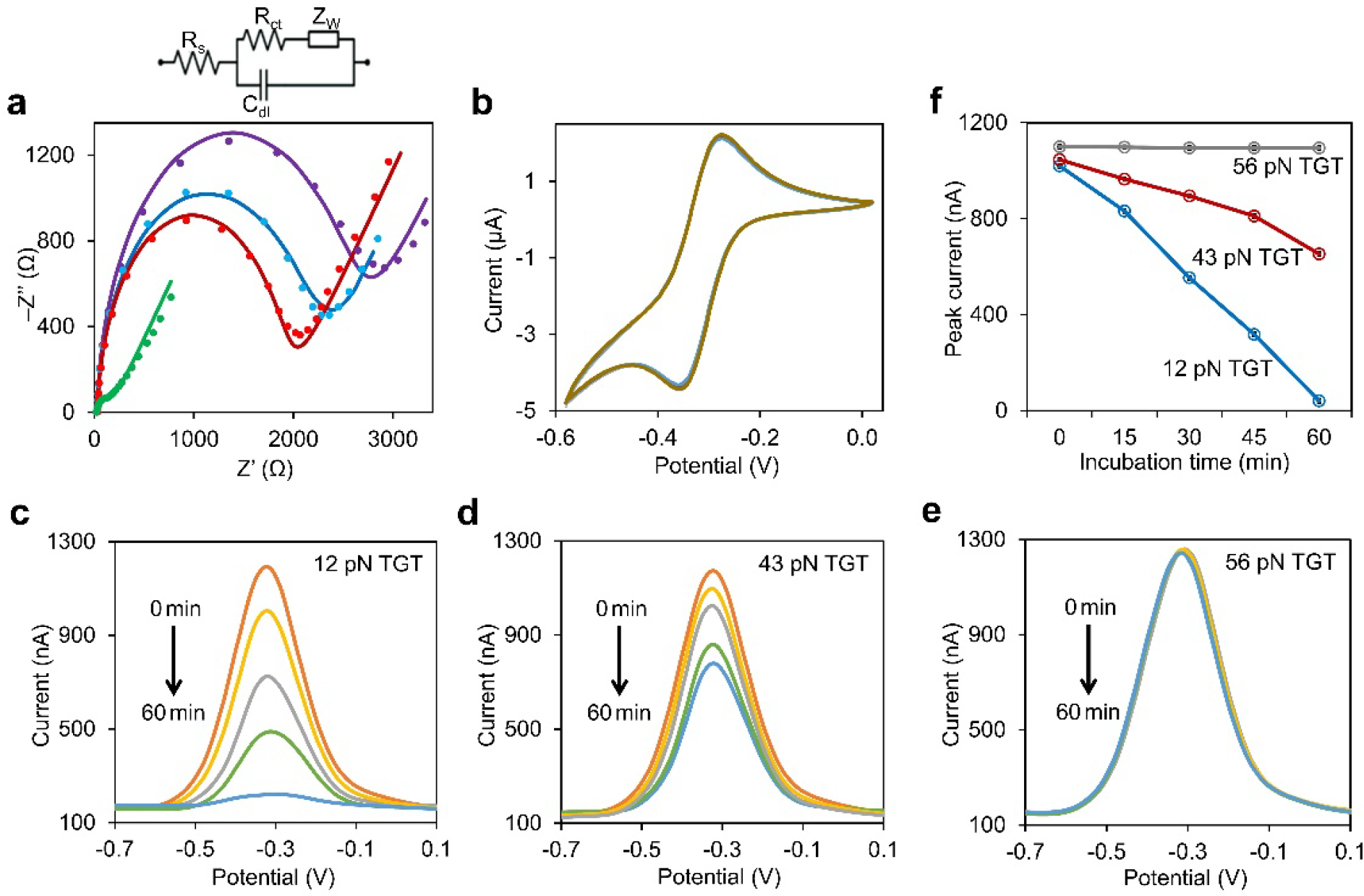
**a**) Nyquist plots of the unmodified Au-SPE (green line), 3’-thiolated anchor strand-modified Au-SPE (red line), 1-hexanethiol-passivated 3’-thiolated anchor strand-modified Au-SPE (blue line), and that after adding 5’-methylene blue-3’-biotin-modified reporter strand (purple line). The electrochemical impedance spectroscopy measurement was performed in a solution containing (v/v) 50% DMEM, 50% phosphate buffer (0.2 M, pH 7.4), and 5 mM ferrocene carboxylate, signals were recorded at an AC potential of 5 mV, a DC potential of 0.17 V, and in the frequency range of 100,000– 0.1 Hz. The equivalent electric circuit compatible with the Nyquist diagrams were shown on the top. Here, *R*_*s*_ is the solution resistance, *R*_*ct*_ is the charge transfer resistance, *C*_dl_ is double layer capacitance, and *Z*_W_ is Warburg impedance. **b**) Stability of the cyclic voltammetry signals of Ru(NH_3_)_6_^3+^ on 1-hexanethiol-passivated thiolated anchor strand-modified Au-SPE during 25 times potential scans at the scan rate of 100 mV.s^−1^ in a solution containing (v/v) 50% DMEM and 50% phosphate buffer (0.2 M, pH 7.4). **c–e**) Square wave voltammetry of the electrochemical **c**) 12 pN TGT sensor, **d**) 43 pN TGT sensor, **e**) 56 pN TGT sensor after adding 100 µL of 1×10^6^ HeLa cells/mL for 0, 15, 30, 45, and 60 min, respectively, followed by washing. The measurement was performed in a solution containing (v/v) 50% DMEM and 50% phosphate buffer (0.2 M, pH 7.4). The step potential was set as 20 mV, the pulse amplitude was at 50 mV, and the frequency was at 20 Hz. **f**) Cell incubation time-dependent changes in the peak current value of 12 pN, 43 pN, and 56 pN TGT sensors. Shown are the mean and standard error peak values after subtracting the background signals from four replicated tests.

We next measured the surface densities of the thiolated anchor strand on the Au-SPE electrode using a Ru(NH_3_)_6_^3+^-based assay.^[15]^ Ru(NH_3_) _6_^3+^ could quantitatively interact with the negatively charged phosphate backbones of DNAs, and then the amount of surface-adsorbed Ru(NH_3_) _6_^3+^ could be determined via the cyclic voltammetry (**CV**). As shown in Figures 2b and S2a, the CV curves of the thiolated anchor strand-modified Au-SPE indeed exhibited a couple of well-defined redox peaks of Ru(NH_3_)_6_^3+^, which were stably shown during 25 times potential scans at the rate of 100 mV·s^-1^. By further increasing the scan rate (**ט**) in the range of 25–300 mV·s^-1^, both anodic and cathodic peak currents (**I**_**p**_) were observed to follow a linear correlation with the changes in the scan rate, indicating a surface-controlled process with the surface coverage of Ru(NH_3_)_6_^3+^ calculated to be ∼4.2×10^−10^ mol·cm^−2^ (Figures S2b and S2c). Correspondingly, the surface coverage of the thiolated anchor strand was estimated to be ∼2.8×10^13^ per cm^2^ (supporting information). The surface of Au-SPE is highly packed with DNA probes.

After the further addition of 5’-MB-3’-biotin-modified reporter strand, the surface coverage of these reporter strands can be directly measured based on the redox peaks of MB in the CV curves (Figure S3a). The CV curves of MB did not exhibit any obvious change after 25 times of scanning the potential in the range of -0.7 V to -0.1 V (Figure S3b), indicating the high stability of the TGT sensor signal for potential precise detection. Similar to the above-mentioned Ru(NH_3_)_6_^3+^-based assay, we further recorded the CV curves of MB at different scan rates (25–300 mV·s^-1^). The peak currents were again linearly proportional to the scan rate (Figure S3c), suggesting a surface-controlled redox process. The surface coverage of MB and consequently the reporter strand were calculated to be ∼2.7×10^13^ per cm^2^. By comparing this value with that of the thiolated anchor strand (i.e., ∼2.8×10^13^ per cm^2^), >95% of the surface-attached anchor strand could be used to generate double-stranded TGT sensors.

### Electrochemical detection of cell-generated adhesion forces with the TGT sensor

To test whether our fabricated TGT sensors can indeed be used to detect cell-generated forces, we first added 100 µL of HeLa cells (1×10^6^ cells/mL concentration) respectively onto the Au-SPE electrodes that were modified with the 12 pN, 43 pN, or 56 pN TGT sensors. HeLa cells were chosen here because their membrane-expressed integrin αvβ3 can specifically recognize the cRGDfK ligand on the TGT sensors and facilitate force-mediated cell adhesion. To study the effect of these adhesion forces, we decided to measure the square wave voltammetry (SWV) signals of the TGT sensors because of the high sensitivity of SWV.^[16]^ As shown in Figure 2c–2e, after a 15–60 min incubation and washing, while almost no change in the peak current signals of the 56 pN TGT sensor was observed, a gradually decreased peak current was shown for both 12 pN and 43 pN TGT sensors.

To further quantify these force-induced SWV signals, we calculated the relative peak current intensity change (I_%_) as I_%_ = (I_c_ – I_0_) / I_0_ ×100%,^[17]^ where I_0_ and I_c_ are the peak current signals of the TGT sensors in the absence and presence of HeLa cells, respectively. After a 60-min incubation with 100 µL of 1×10^6^ HeLa cells/mL, the 12 pN, 43 pN, and 56 pN TGT sensors exhibited respectively an ∼94%, ∼37%, and ∼0.4% decrease in the relative peak current signal (Figure 2f). A decreased SWV peak current suggested a reduced amount of MB-modified reporter strands on the electrode. After incubating with the HeLa cells, the 12 pN and 43 pN TGT sensors could be ruptured and release the reporter strands. In contrast, our results indicate that forces generated by the HeLa cells could not reach 56 pN to rupture the shear-mode TGT sensors.

In the case of the 56 pN TGT sensor, we wondered whether the HeLa cells could indeed adhere to the surface of the electrode, but due to the high threshold force to rupture the TGT probes, the cells will be instead immobilized on the electrode. To test this, we first imaged the surfaces of TGT-modified transparent gold electrodes after a 60-min incubation with HeLa cells and washing. The number of HeLa cells on the surface of the 56 pN TGT electrode was indeed much higher than that of 43 pN and 12 pN sensors (Figure S4a). The EIS and Nyquist plots of the 56 pN TGT sensor were also used to study this cell adhesion process. After incubating with HeLa cells, the R_ct_ value of the sensor increased dramatically from ∼2510 Ω to ∼5670 Ω (Figure S4b), suggesting that less amount of the ferrocene carboxylate redox probe could now reach the surface of the electrode due to mass transfer limitation from the surface-attached HeLa cells. All these data indicated that the HeLa cells could not rapture the 56 pN TGT sensor and thus still attached to the surface of the gold electrode.

We next wanted to test the sensitivity of the electrochemical TGT sensors by detecting adhesion forces from a smaller number of cells. Using the 12 pN TGT sensor as the example, we incubated a total of ∼100, 1×10^3^, 1×10^4^, and 1×10^5^ HeLa cells respectively on the surface of the electrode for 60 min, and afterwards a ∼32%, 57%, 86%, and 94% decrease in the relative SWV peak current intensity was clearly observed (Figure 3a). The TGT sensors could indeed be used to measure forces generated by a small number of cells.

**Figure 3.**
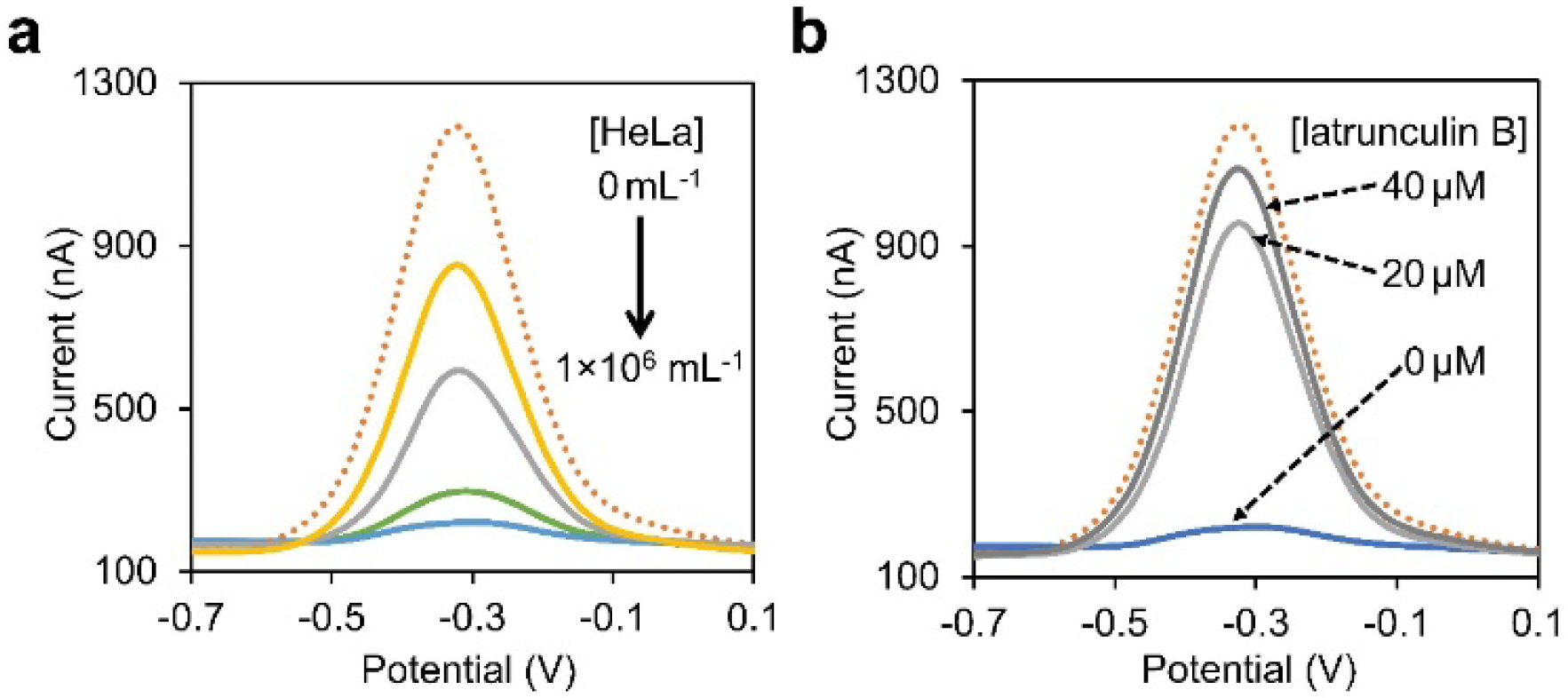
**a**) The square wave voltammetry of the 12 pN TGT sensor after adding 100 µL of 0, 1×10^3^, 1×10^4^, 1×10^5^, or 1×10^6^ HeLa cells/mL for 60 min and washing. The measurement was performed in a solution containing (v/v) 50% DMEM and 50% phosphate buffer (0.2 M, pH 7.4). The step potential was set as 20 mV, the pulse amplitude was at 50 mV, and the frequency was at 20 Hz. Shown are representative curves from four replicated tests. **b**) The square wave voltammetry of the 12 pN TGT sensor after adding 100 µL of 1×10^6^ HeLa cells/mL that have been treated with 0 µM, 20 µM, or 40 µM latrunculin B for 60 min and washing. The measurement was performed in a solution containing (v/v) 50% DMEM and 50% phosphate buffer (0.2 M, pH 7.4). The step potential was set as 20 mV, the pulse amplitude was at 50 mV, and the frequency was at 20 Hz. Shown are representative curves from four replicated tests.

We also wondered whether the TGT sensors could also be applied to study the effect of potential force-inhibiting drugs on the cell-generated forces. For this purpose, we chose to test latrunculin B, an inhibitor of actin polymerization that can bind to cellular G-actin and prevent its transition into F-actin.^[18]^ As shown in Figure 3b, in the presence of 20 µM or 40 µM of latrunculin B, after incubating 100 µL of 1×10^6^ HeLa cells/mL with the 12 pN TGT electrode for 60 min, the I_%_ value of the sensor was significantly changed to ∼22% and 9%, as compared to that of ∼94% in the absence of force-inhibiting drugs. As expected, after the inhibition of actin polymerization, the integrin αvβ3 of the HeLa cells could not apply the forces to rupture the TGT sensors. All these above-shown results demonstrated that our proposed electrochemical TGT sensors can indeed be used as a highly sensitive and easy-to-use tool for detecting cell-generated adhesion forces.

### Construction and characterization of the electrochemical DNA hairpin-based force sensors

Our next goal was to study whether electrochemical DNA hairpin-based sensors could be developed to measure cellular adhesion forces. In our design, the hairpin force sensor comprises a 5’-RGD-3’-thiol-modified anchor hairpin strand and a 5’-MB-modified reporter strand (Table S1). Once the cRGDfK ligands experience sufficient tensions (in this case, ∼18 pN) from cellular integrin αvβ3, the unfolding of DNA hairpins will result in an increased distance of MB from the Au-SPE surface, and consequently, a decreased electrochemical signal.^[19]^ In contrast to the TGT sensors, hairpin-based force sensors can be more reversibly open and close to monitor dynamic changes in the cellular forces.

The construction of hairpin force sensors started by annealing a 5’-biotin-3’-thiol-modified anchor strand with a 5’-MB-labeled reporter strand, which duplex was then added and self-immobilized onto surface-cleaned Au-SPE. Like the TGT force sensors, after washing, 1-hexanethiol passivation, and stepwise addition of streptavidin and biotinylated cRGDfK, the DNA hairpin probes were finally fabricated on the surface of the Au-SPE electrode (Figure 1).

EIS was also used to characterize these construction steps. As shown in Figure 4a, the unmodified Au-SPE exhibited a low R_ct_ at ∼125 Ω, while after adding the conjugates of thiolated anchor strand and MB-labeled reporter strand, the R_ct_ value was increased to ∼1270 Ω, indicating a layer of DNA conjugates was immobilized on the surface of the electrode that could electrostatically repulse the negatively charged ferrocene carboxylate. Based on these R_ct_ values, the surface coverage of the DNA strand θ_(t)_ was found to be ∼90%. After additionally assembling streptavidin and biotinylated cRGDfK, the R_ct_ value increased to ∼2140 Ω, suggesting a further limited mass transfer of ferrocene carboxylate to the electrode surface.

**Figure 4.**
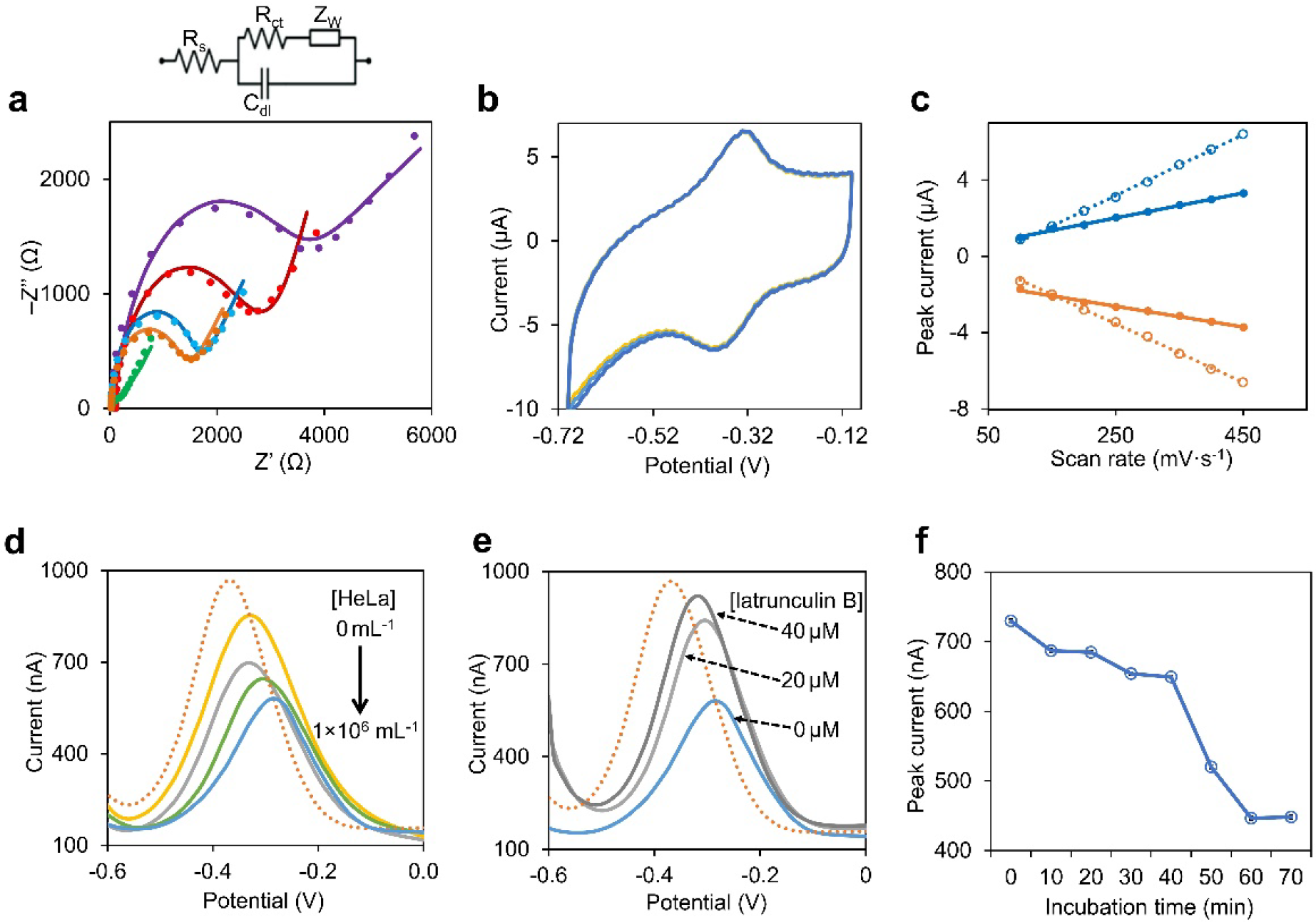
**a**) Nyquist plots of the unmodified Au-SPE (green line), thiolated DNA hairpin probe-modified Au-SPE (orange line), 1-hexanethiol-passivated thiolated DNA hairpin probe-modified Au-SPE (blue line), 1-hexanethiol-passivated thiolated DNA hairpin probe-modified Au-SPE after adding streptavidin and biotinylated cRGDfK, i.e., DNA hairpin force sensor (red line), and that after adding 100 µL of 1×10^6^ HeLa cells/mL onto the DNA hairpin force sensor (purple line). The electrochemical impedance spectroscopy measurement was performed in a solution containing (v/v) 50% DMEM, 50% phosphate buffer (0.2 M, pH 7.4), and 5 mM ferrocene carboxylate, signals were recorded at an AC potential of 5 mV, a DC potential of 0.17 V, and in the frequency range of 100,000–0.1 Hz. The equivalent electric circuit compatible with the Nyquist diagrams were shown on the right. *R*_*s*_ is the solution resistance, *R*_*ct*_ is the charge transfer resistance, *C*_dl_ is double layer capacitance, and *Z*_W_ is Warburg impedance. **b**) Stability of the cyclic voltammetry signals of methylene blue on the DNA hairpin force sensor-modified Au-SPE during 25 times potential scans at the scan rate of 400 mV.s^−1^ in a solution containing (v/v) 50% DMEM and 50% phosphate buffer (0.2 M, pH 7.4). **c**) Before (dotted line) and after (solid line) adding HeLa cells, a linear correlation was also observed between both anodic (orange line) and cathodic (blue line) peak currents (I_p_) and the changes in the scan rate (ט). **d**) The square wave voltammetry of the DNA hairpin sensor after adding 100 µL of 0, 1×10^3^, 1×10^4^, 1×10^5^, or 1×10^6^ HeLa cells/mL for 60 min. The measurement was performed in a solution containing (v/v) 50% DMEM and 50% phosphate buffer (0.2 M, pH 7.4). The step potential was set as 20 mV, the pulse amplitude was at 50 mV, and the frequency was at 20 Hz. Shown are representative curves from four replicated tests. **e**) The square wave voltammetry of the DNA hairpin sensor after adding 100 µL of 1×10^6^ HeLa cells/mL that have been treated with 0 µM, 20 µM, or 40 µM latrunculin B for 60 min. The measurement was performed in a solution containing (v/v) 50% DMEM and 50% phosphate buffer (0.2 M, pH 7.4). The step potential was set as 20 mV, the pulse amplitude was at 50 mV, and the frequency was at 20 Hz. Shown are representative curves from four replicated tests. **f**) Cell incubation time-dependent changes in the peak current value of the DNA hairpin sensor. Shown are the mean and standard error peak values after subtracting the background signals from four replicated tests.

To determine the surface coverage of the hairpin probes, we next measured the CV curves of MB on the Au-SPE electrode at a scan rate of 400 mV·s^-1^ (Figure S5a). The paired redox peaks of MB were stably shown during 25 times of potential scans (Figure 4b), indicating a high stability of the electrochemical DNA hairpin-based sensor. By recording the CV curves at different scan rates in the range of 100–450 mV·s^-1^, both the oxidation and reduction peaks of MB followed a linear relationship with the scan rate (Figures 4c and S5b). Similar to the TGT sensors, the surface coverage of DNA hairpin probes can be calculated as a surface-controlled electrochemical system, with ∼1.9×10^13^ probes being modified per cm^2^ electrode (supporting information).

### Electrochemical detection of cell-generated adhesion forces with the DNA hairpin sensor

We next wanted to study whether the surface attachment of HeLa cells will induce the opening of DNA hairpin probes. For this purpose, we measured the heterogeneous electron transfer rate constant (**K**_**s**_) of MB, as the K_s_ value highly depends on the distance of MB from the surface of Au-SPE.^[19]^ Upon the unfolding of DNA hairpin, a dramatically decreased K_s_ will be observed. After adding 100 µL of HeLa cells (1×10^6^ cells/mL) for 60 min, the CV curves of the hairpin sensor were recorded at different scan rates (100–450 mV·s^-1^). Based on a Laviron’s formula,^[20]^ the average K_s_ of MB was calculated to be ∼0.21 s^-1^ (Figures 4c and S5c). While in the absence of cells, a K_s_ value of ∼7.1 s^−1^ was shown. This dramatic decrease in the K_s_ of MB (∼33-fold) indicated that the distance between MB and the surface of the electrode was indeed significantly increased after the adhesion of HeLa cells.

To further validate whether the surface attachment of HeLa cells indeed depends on the specific interactions between integrin αvβ3-RGD and cRGDfK, EIS measurements were performed to measure cell adhesion-induced changes in the charge-transfer resistance of the electrode. As expected, after a 60-min incubation with HeLa cells (100 µL of 1×10^6^ cells/mL), the R_ct_ of the hairpin sensor increased dramatically from ∼2140 Ω to ∼3420 Ω (Figure 4a), indicating that the surface attachment of HeLa cells hindered ferrocene carboxylate from reaching the surface of the electrode. While in contrast, after the same batch of HeLa cells was incubated with a control Au-SPE electrode that was immobilized with the same DNA hairpin probe just without the attachment of biotinylated cRGDfK, a minimal change in the R_ct_ value (from ∼1590 Ω to ∼1650 Ω) supported that the electrode surface adhesion of HeLa cells highly relies on the presence of integrin–RGD interactions (Figure S6a).

Our next goal was to study the sensitivity of the hairpin sensor in measuring cell-generated adhesion forces. A total of ∼100, 1×10^3^, 1×10^4^, and 1×10^5^ HeLa cells were added and incubated respectively for 60 min on the surface of the hairpin sensor. As shown in Figure 4d, the SWV peak current of the electrode continuously decreased as the concentration of cells increased, suggesting that more and more DNA hairpins were unfolded by surface-attached HeLa cells. We also calculated the relative peak current intensity change of the hairpin sensors. Our results showed that the I_%_ values changed from ∼7%, 25%, 33%, to ∼42% as the amount of HeLa cells increased from 100 to 1×10^5^. Like the TGT sensors, hairpin-based electrochemical sensors could also be used to measure forces generated from a small number of cells.

As another control experiment to test if the observed changes in the SWV peak current could be possibly due to simple cell attachment but without the opening of DNA hairpins, we also immobilized cRGDfK ligands directly onto the surface of Au-SPE, together with cRGDfK-free DNA hairpin probes. In the presence of ferrocene carboxylate, we then measured the SWV signals of both MB and ferrocene carboxylate before and after a 60-min incubation with 1×10^3^ HeLa cells (Figure S6b). The peak current of ferrocene carboxylate decreased significantly due to mass transfer limitation caused by the attachment of HeLa cells. While in contrast, the MB signals exhibited no change after adding the HeLa cells. These results indicated that without integrin–RGD interaction-mediated unfolding of DNA hairpins, simple attachment of HeLa cells will not affect the hairpin sensor signals.

The hairpin force sensor can also be used to detect the force-inhibition function of latrunculin B. After adding 20 µM or 40 µM of latrunculin B to 1×10^5^ HeLa cells and incubating for 60 min, the SWV peak current of the hairpin sensor increased dramatically (with an I_%_ value of ∼19% and 8%) as compared to that without the drug treatment (∼42% I_%_) (Figures 4e). This result was expected, as the inhibition of integrin forces could result in less amount of DNA hairpin probes being unfolded.

Lastly, we wondered whether the hairpin force sensor could be applied to monitor the dynamic changes in the cell-generated adhesion forces. To test this, the SWV response of the sensor was measured every 10 min after adding 1×10^5^ HeLa cells. Interestingly, our results indicated that during the first 40-min incubation, a slow decrease in the SWV peak current was observed, while afterward, a much faster and more dramatic change was shown (Figures 4f). Since the initial HeLa cell surface attachment occurred immediately after adding the cells onto the electrode, we realize that the observed 40-min delay in the SWV signal change is likely due to the time requirement for cell spreading and focal adhesion formation.^[11b]^ Upon forming focal adhesions, integrins could then apply strong forces to the cRGDfK ligand and open the DNA hairpin probe, which has a force threshold value of ∼18 pN. Again, all these data suggested the power and feasibility of electrochemical DNA hairpin-based sensor for measuring cellular adhesion forces.

## Discussions

In this project, three types of TGT probes and also a DNA hairpin probe have been fabricated onto a smartphone-based electrochemical device for the sensitive and robust detection of cell-generated adhesion forces. By simply changing the DNA sequence or force application geometry, these electrochemical DNA-based sensors could modularly detect cellular forces at different threshold values. Meanwhile, the in situ dynamic changes in cellular adhesion forces as well as the regulation effect of force inhibitor drugs could also be measured using these novel electrochemical sensors. In addition, by potentially altering the ligand modification of the DNA probes, cellular adhesions mediated by different membrane ligand–receptor pairs could be possibly investigated at the molecular level.

As far as we know, this is the first attempt to measure cellular forces using electrochemical DNA-based molecular sensors. Compared to previously reported fluorescence-based force sensors,^[10,11]^ these electrochemical tools could be more advantageous in terms of their portability, cost efficiency, and simplicity of use. We expect these sensors could be potentially useful for studying fundamental cell signaling pathways and also for developing novel strategies and materials in tissue engineering and pharmaceutics.

## Data availability

The data supporting the findings of this study are available upon request to the corresponding author.

## Acknowledgements

The authors gratefully acknowledge the support from NIH R35GM133507, T32GM139789, and Camille Dreyfus Teacher-Scholar Award. The authors also thank other members of the You Lab for useful discussion and valuable comments.

## Additional information

### Supplementary information

The online version contains supplementary material available.

