## Supplementary Information for "Electrochemical DNA-based sensors for measuring cell-generated forces"

### 1. Materials and Methods

**Reagents and apparatus.** Double deionized water (18.6 MΩ·cm$^{-1}$) was used throughout the research work. The DNA oligonucleotides were custom synthesized and purified by W. M. Keck Oligonucleotide Synthesis Facility at Yale University School of Medicine. Unless otherwise mentioned, all the chemicals were purchased from Thermo Fisher Scientific (Agawam, MA) and used without further purification. Gold screen-printed electrodes (Au-SPE) and transparent gold electrodes were purchased from Metrohm-DropSens (Llanera, Spain). The cyclic voltammetry, electrochemical impedance spectroscopy, and square wave voltammetry studies were performed using an electrochemical device from PalmSens Model Sensit Smart (Houten, Netherlands).

**Cleaning the surface of the electrodes.** The surface of gold electrodes was cleaned in 2.0 M $H_2SO_4$ solution. After the cleaning, by scanning the potential between -0.3 V and 1.2 V, a sharp cathodic peak at ~0.56 V related to the electrochemical reduction of Au oxide to metallic Au could be observed. Cleaning was performed until there was no further change in the cyclic voltammogram. Meanwhile, the roughness factor (RF) of the cleaned electrode surface can be calculated as $A_{eas}/A_{gms}$, where $A_{eas}$ is the electroactive surface area and $A_{gms}$ is the geometric surface area. $A_{eas} = Q_{Faradic}$ / (390 μC·cm$^{-2}$).[1] Here, 390 μC·cm$^{-2}$ is the conversion factor of the surface area of polycrystalline gold, and $Q_{Faradic}$ is the faradic charge associated with the reduction of Au oxide that can be determined by integrating the reduction peak current area (between -0.40 V and -0.75 V) and then divided by the scan rate (10.0 mV·s$^{-1}$). As shown in Figure S1a, in this study, $A_{gms}$ was ~0.125 cm$^2$, $Q_{Faradic}$ was found to be ~49.9 μC, and $A_{eas}$ was ~0.127 cm$^2$. A ~1 roughness factor suggesting a well-cleaned surface of gold electrodes.

**Preparation of electrochemical TGT sensors.** The thiolated anchor strand (0.05 mM) was first reduced using 10 mM TCEP for 1 h to cleave the disulfide bonds. This solution was then diluted by 0.1 M phosphate buffer (pH 7.4) to obtain 5.0 μM thiolated DNA. Afterwards, 100 μL of 5.0 μM thiolated anchor strand was dropped onto the cleaned Au-SPE surface and kept in the refrigerator for 16 h. During this incubation time, the thiolated anchor strand bonded to the Au-SPE surface by self-assembly through the Au–S bonds.[2] After rinsing with copious amounts of 0.1 M phosphate buffer (pH 7.4) to wash away nonspecifically adsorbed DNA molecules, 100 μL of 100 μM 1-hexanethiol solution was added on the surface of the electrode and incubated at 37°C for 1 h to passivate the remaining active sites of the electrode surface. Subsequently, 100 μL of 5.0 μM 5'-MB-3'-biotin-modified reporter strand was added to the surface of the electrode and incubated at room temperature for 1 h to generate surface-attached double-stranded DNA. After washing profusely with 0.1 M phosphate buffer (pH 7.4), 100 μL of 5.0 μM streptavidin (in 0.1 M phosphate buffer, pH 7.4) was dropped on the surface of the electrode to interact with the biotin group of DNA duplex at room temperature for 1 h. After again rinsing plentifully with 0.1 M phosphate buffer (pH 7.4), 100 μL of 5.0 μM biotinylated cyclic arginine-glycine-aspartic-acid-D-phenylalanine-lysine (cRGDfK) was dropped on the surface of the electrode to interact with streptavidin at room temperature for 1 h. As a final step, the fabricated TGT sensor was rinsed with 0.1 M phosphate buffer (pH 7.4) and stored at 4°C before usage.

**Preparation of electrochemical DNA hairpin sensors.** 100 μL of 50 μM 5'-biotin-3'-thiol-modified hairpin anchor strand was first mixed with 100 μL of 5'-methylene blue (MB)-labeled reporter strand for 15 min, and then the mixture was annealed by heating to 75°C for 5 min followed by cooling to 25°C at a rate of –1.3°C/min using a Bio-Rad PCR thermal cycler. During this annealing procedure, two DNA strands hybridized with each other. Then following a similar process as the TGT sensors, the DNA duplex was reduced using 10 mM TCEP for 1 h to cleave the disulfide bonds. This solution was then diluted by 0.1 M phosphate buffer (pH 7.4) to obtain 5.0 μM DNA. Afterwards, 100 μL of 5.0 μM thiolated hairpin anchor strand was dropped onto the cleaned Au-SPE surface and kept in the refrigerator for 16 h. During this incubation time, the thiolated anchor strand bonded to the Au-SPE surface by self-assembly through the Au–S bonds.[2] After rinsing with copious amounts of 0.1 M phosphate buffer (pH 7.4) to wash away nonspecifically adsorbed DNA molecules, 100 μL of 100 μM 1-hexanethiol solution was added on the surface of the electrode and incubated at 37°C for 1 h to passivate the remaining active sites of the electrode surface. Subsequently, 100 μL of 5.0 μM streptavidin (in 0.1 M phosphate buffer, pH 7.4) was dropped on the surface of the electrode to interact with the biotin group of DNA probes at room temperature for 1 h. After rinsing plentifully with 0.1 M phosphate buffer (pH 7.4), 100 μL of 5.0 μM biotinylated cyclic arginine-glycine-aspartic-acid-D-phenylalanine-lysine (cRGDfK) was dropped on the surface of the electrode to interact with streptavidin at room temperature for 1 h. As a final step, the fabricated DNA hairpin sensor was rinsed with 0.1 M phosphate buffer (pH 7.4) and stored at 4°C before usage.

**Determining the surface coverage of DNA probes.** After thiolated anchor strand was immobilized on the surface of gold electrode and passivated with 1-hexanethiol, 100 μL of 1 mM hexaammineruthenium (III) ion ($Ru(NH_3)_6^{3+}$) was dropped on the electrode surface and incubated for 1 h at room temperature. During this time, the positively charged $Ru(NH_3)_6^{3+}$ were assembled with the negatively charged phosphate backbone of DNA.[3] After washing with 0.1 M phosphate buffer (pH 7.4), cyclic voltammetry was performed by increasing the scan rate (ʋ) in the range of 25–300 $mV\cdot s^{-1}$. In this case, both anodic and cathodic peak currents ($I_p$) were observed to follow a linear change with the scan rate, indicating a surface-controlled process so that the surface coverage of $Ru(NH_3)_6^{3+}$ (Γ) could be calculated based on $I_p = (\Gamma\cdot A\cdot n^2\cdot F^2)\cdot \upsilon /(4R\cdot T)$,[4] where n is the number of electrons released ($Ru^{3+}/Ru^{2+}$, n=1), A is the electroactive surface area (0.127 $cm^2$ in this case), F is the Faraday constant (96,485 $C.mol^{-1}$), R is the universal gas constant (8.314 $J\cdot K^{-1}\cdot mol^{-1}$), and T is the temperature (298.15 K). By analyzing the slope of the $I_p$ versus scan rate (ʋ), the average $\Gamma_{Ru}$ could be calculated to be ~$4.2\times10^{-10}$ $mol\cdot cm^{-2}$. Based on this, the surface coverage of the thiolated anchor strand on the gold electrode could be calculated by using the equation[5] $\Gamma_{Anchor\ DNA} = \Gamma_{Ru}\cdot z\cdot N_A/m$, where z is the charge of $Ru(NH_3)_6^{3+}$ (3), m is the number of $Ru(NH_3)_6^{3+}$ that could interact with each thiolated anchor strand (21), and $N_A$ is the Avogadro's number ($6.022\times10^{23}$ $mol^{-1}$). In our case, the $\Gamma_{Anchor\ DNA}$ was calculated to be ~$2.8\times10^{13}$ per $cm^2$.

The surface coverage of 5'-MB-3'-biotin-modified reporter strand was determined using a similar method but was based on the cyclic voltammetry redox peaks of MB rather than that of $Ru(NH_3)_6^{3+}$. The number of electrons released (n) is 2 for an MB probe, and since the charge (z) of MB is 1 and the number of MB in each reporter strand (m) is 1, the surface coverage of MB is equal to that of the reporter strand. The $\Gamma_{Reporter\ DNA}$ was calculated to be ~$2.7\times10^{13}$ per $cm^2$. Similar to the reporter strand of TGT sensors, the surface coverage of DNA hairpin probes can also be calculated based on MB cyclic voltammetry signals. The $\Gamma_{Hairpin\ DNA}$ was calculated to be ~$1.9\times10^{13}$ per $cm^2$.

**Sensing process of cell-generated forces.** HeLa cells were cultured in DMEM with 10% fetal bovine serum, 100 U/mL penicillin, and 100 U/mL streptomycin in an Eppendorf Galaxy incubator at 5% (v/v) $CO_2$. Before measurement, Hela cells were first detached by adding 2 mM EDTA in 0.1 M HEPES buffer (pH 7.6) for 10 min,[6] The solution was then centrifuged three times at 1200 rpm for 7 min. After that, the cells were re-suspended in a measuring solution containing (v/v) 50% DMEM and 50% phosphate buffer (0.2 M, pH 7.4), with a final cell concentration from $1\times10^3$ cells/mL to $1\times10^6$ cells/mL. To measure HeLa cell-generated forces, 100 μL of this cell solution was dropped onto the surface of DNA probe-modified gold electrode. The whole sensor device was then put inside a small box and kept inside an Eppendorf Galaxy cell culture incubator. The signal of the electrode was recorded every 10 min using square wave voltammetry, with the step potential set as 20 mV, pulse amplitude at 50 mV, and frequency at 20 Hz. The designed electrochemical force sensors were attached to a smartphone-controlled potentiostat device, which together function as a portable, cost-effective, and sensitive tool for monitoring the forces generated by live cells.

**Heterogeneous electron transfer rate constant ($K_s$) measurement.** To measure $K_s$, the CV curves of the DNA hairpin sensor were recorded at different scan rates in the range of 100–450 $mV\cdot s^{-1}$. For a surface-controlled electrochemical system, in the absence of cells, considering the difference between the anodic and cathodic peak potentials, $\Delta E_p < 200/n$ mV, and also the charge transfer coefficient, $\alpha = 0.5$, a Laviron's formula[7] should be used to calculate the corresponding $K_s$ value: $K_s = (m \cdot n \cdot F \cdot \upsilon)/(R \cdot T)$, where m is the parameter related to the peak potential separation, n is the number of electrons involved in the reaction, $\upsilon$ is the scan rate ($V \cdot s^{-1}$), F is the Faraday constant (96,485 $C \cdot mol^{-1}$), R is the universal gas constant (8.314 $J \cdot K^{-1} \cdot mol^{-1}$), and T is the temperature (298.15 K). In the absence of HeLa cells, the $K_s$ value of the sensor was found to be ~7.1 $s^{-1}$.

After adding 100 µL of HeLa cells ($1 \times 10^6$ cells/mL) for 60 min, the CV curves of the hairpin sensor changed dramatically (Figure S5). Considering the peak-to-peak separation ($\Delta E_p$) of the CV curves became greater than 200/n mV, a revised Laviron's formula should be used for calculating $K_s$ in the presence of HeLa cells, i.e., $\log K_s = \alpha \cdot \log(1 - \alpha) + (1 - \alpha) \cdot \log\alpha - \log[R \cdot T/(n \cdot F \cdot \upsilon)] - \alpha \cdot (1-\alpha) \cdot n \cdot F \cdot \Delta E_p / (2.3 R \cdot T)$.[7] In this case, $\alpha = 0.37$, which was obtained by analyzing the relationship of the reduction potential, $E_{pc}$, versus the logarithm of the scan rate ($\upsilon$) as $E_{pc} = E^0 - 2.3 R \cdot T \cdot \log\upsilon / (\alpha \cdot n \cdot F)$.[7] Based on that, the average $K_s$ of the hairpin sensor after incubating with the HeLa cells was calculated to be ~0.21 $s^{-1}$.

**Force inhibitor treatment.** To study the impact of force inhibition drug on the cellular force generations, before plating the HeLa cells, 0.8 µL or 1.6 µL of 25 mM of latrunculin B was added to 1 mL mixture of (v/v) 50% DMEM and 50% 0.2 M phosphate buffer (0.2 M, pH 7.4) containing $1 \times 10^6$ HeLa cells/mL and kept inside a cell culture incubator for 1 h. After that, 100 µL of the solution was dropped on the surface of either the TGT sensor or DNA hairpin sensor and kept inside a cell culture incubator for another 1 hour, before measuring the electrochemical signal.

**Rapture force estimation of TGT probes.** The irreversible rupture of TGT depends on the orientation of the applied forces and also the length of the duplex region. The tension tolerance ($T_{tol}$) of TGT probes is defined as the rupture force needed to unfold 50% of the DNA duplex. Based on a de Gennes model,[8] $T_{tol}$ can be calculated as $T_{tol} = 2F_c \cdot [X^{-1} \cdot \tanh(X \cdot L/2) + 1]$, where $F_c$ is the rupture force per base pair (3.9 pN), L is the number of base pairs between the points of force application on the two complementary strands of TGT probes, and X is a function describing the elasticity within the dsDNA, dictated by a spring constant (Q) between neighbors within a strand and a spring constant (R) between base pairs in a duplex: $X = (2R/Q)^{1/2}$. Assuming a constant force is applied for 1–2 s, then $X^{-1} = 6.8$.

**Unfolding force estimation of DNA hairpin probes.** The mechanical stability of the hairpin can be analyzed by the $F_{1/2}$ value, i.e., the force at which 50% of hairpins possibly being unfolded. $F_{1/2}$ can be calculated based on the Woodside equation[9] as $F_{1/2} = (\Delta G_{fold} + \Delta G_{stretch})/\Delta x$, where $\Delta G_{fold}$ is the free energy to unfold the DNA hairpin in the absence of forces, $\Delta G_{stretch}$ is the free energy to stretch the unfolded structure of the hairpin from no force to $F = F_{1/2}$, and $\Delta x$ is the hairpin extension length or the opening distance of the hairpin during unfolding. $\Delta G_{stretch}$ can be calculated by $\Delta G_{stretch} = (k_B \cdot T \cdot L_0/L_p) \cdot (3x^2/L_0^2 - 2x^3/L_0^3) / [4 \cdot (1 - x/L_0)]$, where $k_B$ is the Boltzmann constant, T is temperature, $L_p$ is the persistence length of single-stranded DNA (~1.3 nm), $L_0$ is the contour length of single-stranded DNA (~0.63 nm per nucleotide), and x is the hairpin extension from equilibrium, i.e., $0.44 \cdot (n - 1)$ nm, here n is the number of nucleotides in the hairpin. The hairpin extension length $\Delta x$ can be similarly calculated as $[0.44 \cdot (n - 1) - 2]$ nm, 2 nm is subtracted by considering the diameter of the hairpin stem duplex as the initial distance between the hairpin termini.

### 2. Supplementary Table

**Table S1.** The sequence of oligonucleotides used in this study.

| Name | DNA sequence (5’ - 3’) |
|---|---|
| TGT reporter strand | Methylene blue-CACAGCACGGAGGCACGACAC-Biotin |
| 12 pN anchor strand | Thiol-$C_6$-GTGTCGTGCCTCCGTGCTGTG |
| 43 pN anchor strand | GTGTCGTGCCT-Thiol-$C_6$-CCGTGCTGTG |
| 56 pN anchor strand | GTGTCGTGCCTCCGTGCTGTG-$C_6$-Thiol |
| Hairpin reporter strand | Methylene blue-GCTGGGCTACGTGGCGCTCTT |
| Hairpin anchor strand | Biotin-AAGAGCGCCACGTAGCCCAGCGCGCGCGCGCTTTTGCG CGCGCGCGC-$C_6$-Thiol |

### 3. Supplementary Figures

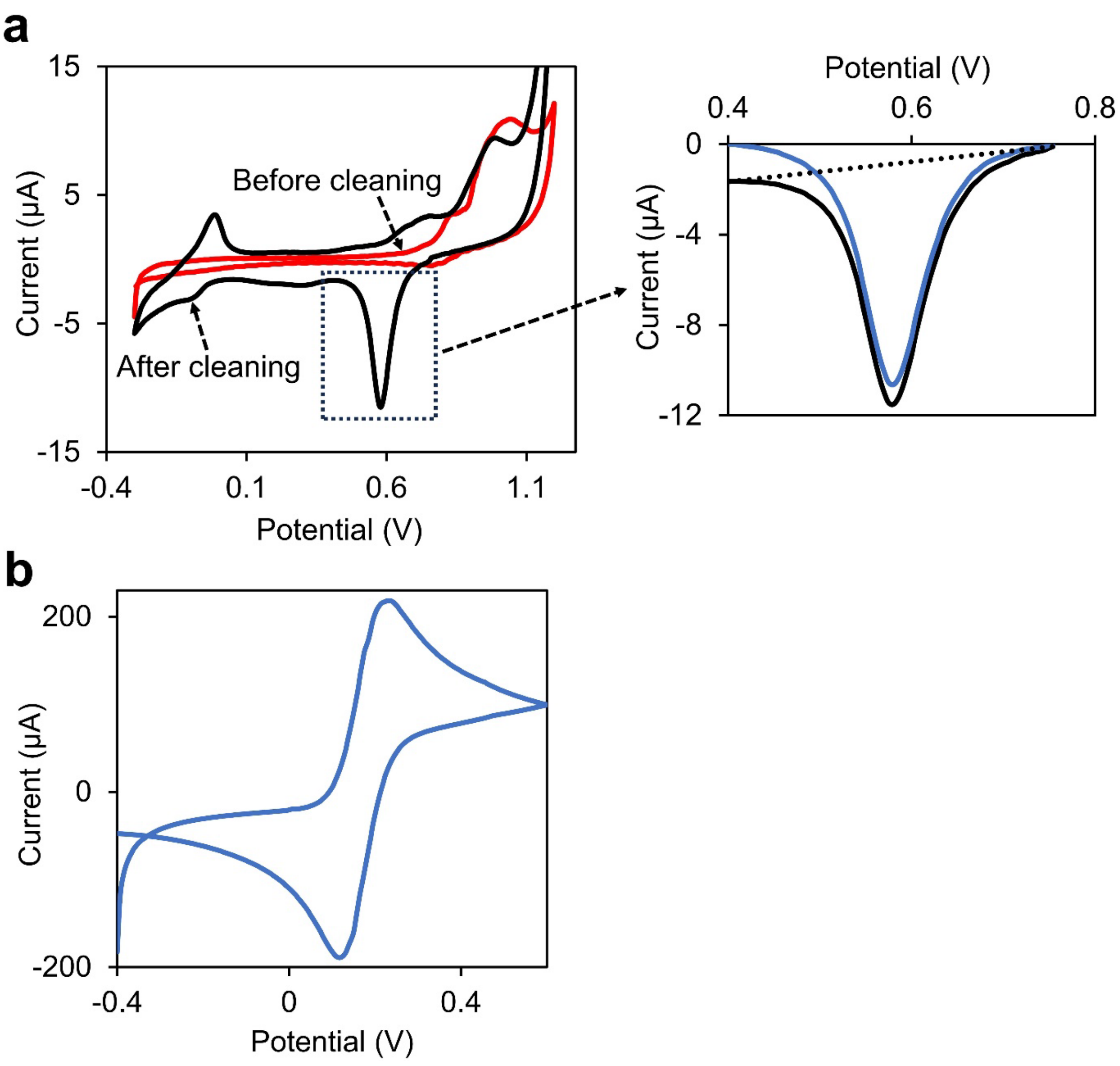


**Figure S1.** (**a**) Cyclic voltammogram of the Au-SPE before (red line) and after (black line) cleaning with 2.0 M $H_2SO_4$ solution. The scan rate was at 10.0 mV·s$^{-1}$. A sharp cathodic peak at ~0.56 V was related to the electrochemical reduction of Au oxide to metallic Au. (Right) A zoomed-in view of this cathodic peak and the baseline adjustment (blue line) for the calculation of peak area. (**b**) Cyclic voltammogram of 5 mM ferrocene carboxylate in a solution containing (v/v) 50% DMEM, 50% phosphate buffer (0.2 M, pH 7.4) at a scan rate of 100 mV·s$^{-1}$.

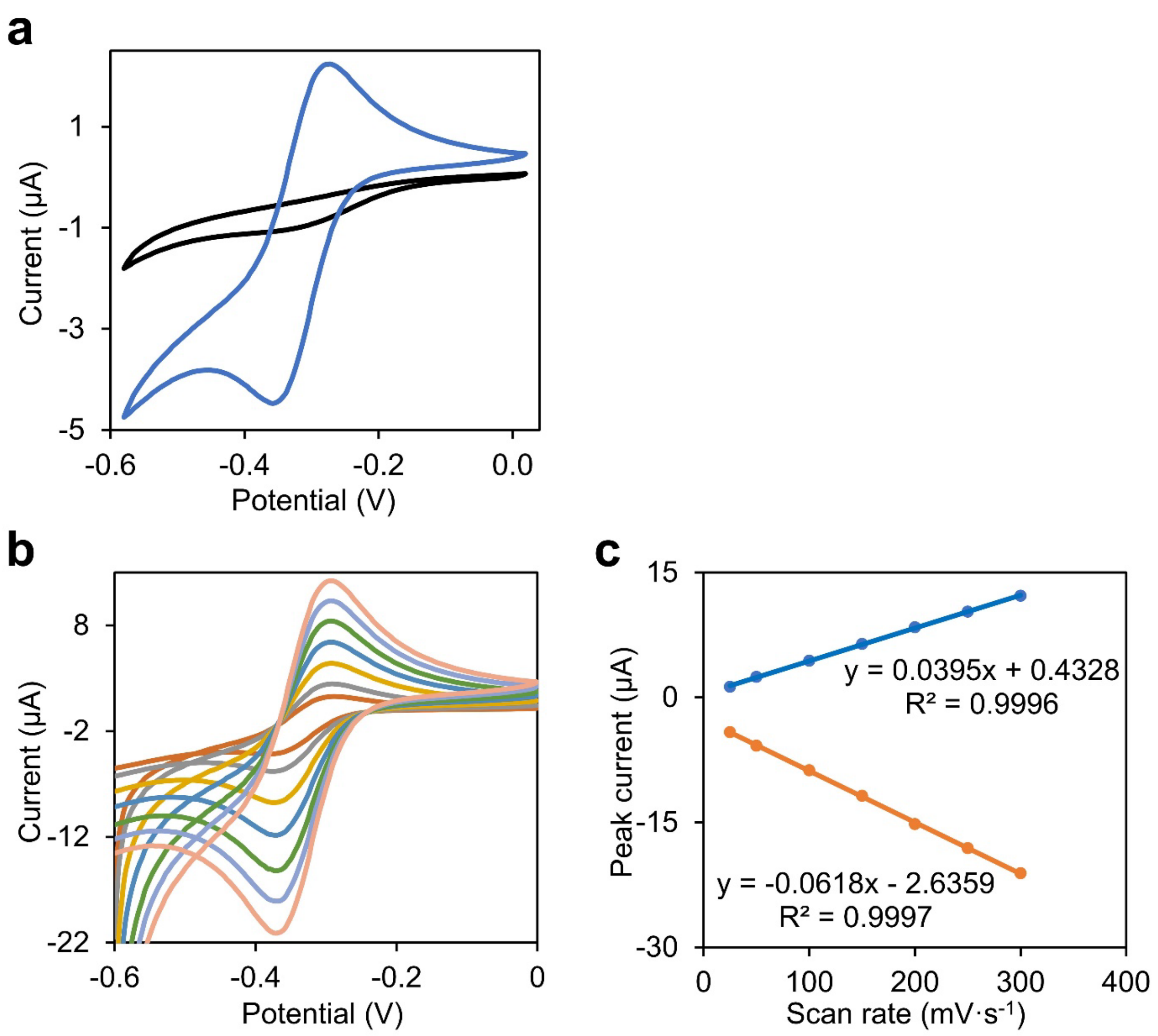


**Figure S2.** (**a**) Cyclic voltammogram of 1-hexanethiol-passivated thiolated anchor strand-modified Au-SPE before (black line) and after (blue line) interaction with 5.0 μM $Ru(NH_3)_6^{3+}$ in a solution containing (v/v) 50% DMEM and 50% phosphate buffer (0.2 M, pH 7.4) at a scan rate of 100 $mV{\cdot}s^{-1}$. (**b**) Cyclic voltammogram of $Ru(NH_3)_6^{3+}$ on 1-hexanethiol-passivated thiolated anchor strand-modified Au-SPE at various scan rates of 25, 50, 100, 150, 200, 250, and 300 $mV{\cdot}s^{-1}$ from inner to outer. (**c**) A linear correlation was observed between both anodic (orange line) and cathodic (blue line) peak currents ($I_p$) and the changes in the scan rate ($\upsilon$).

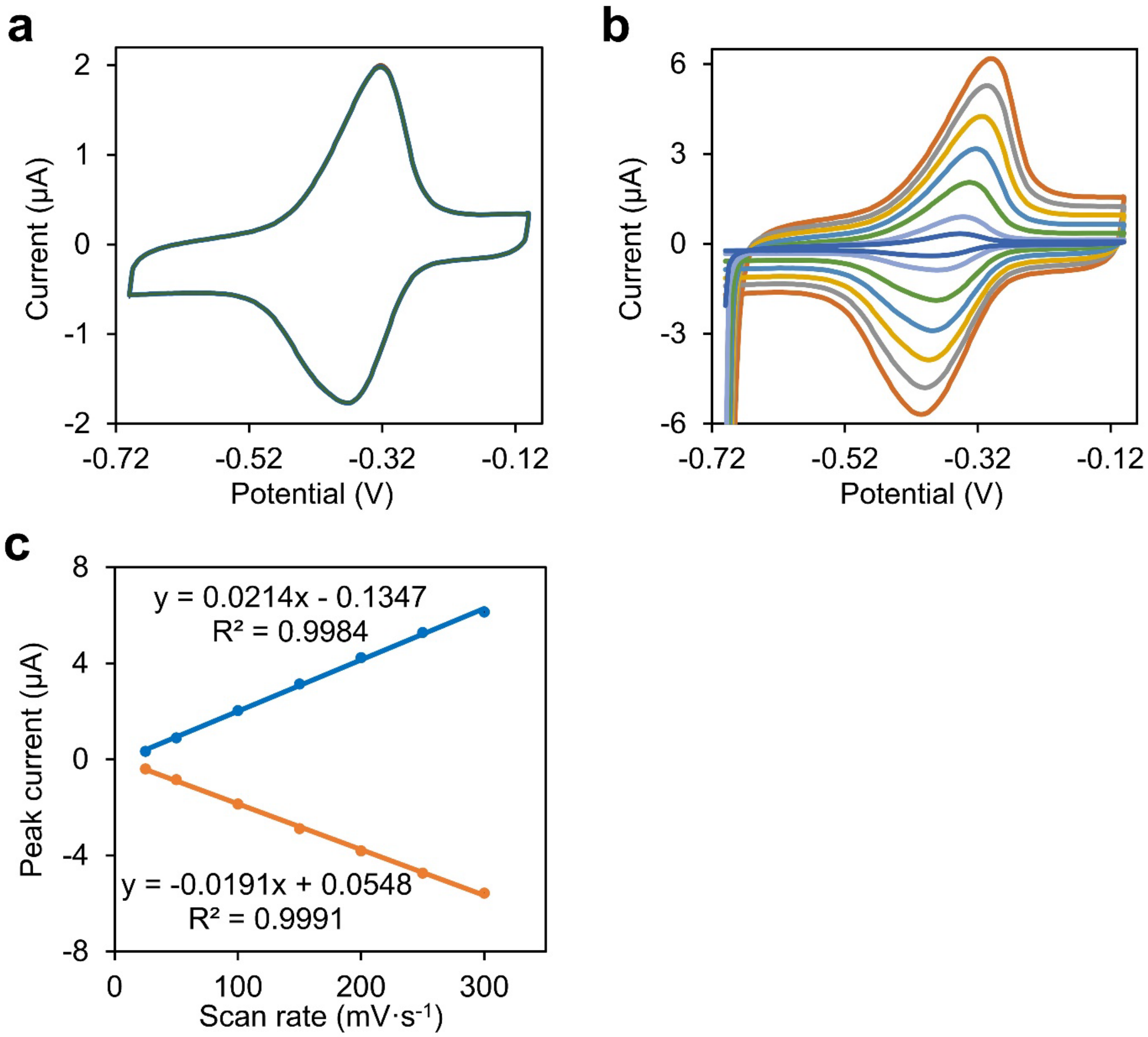


**Figure S3.** (**a**) Stability of the cyclic voltammetry signals of methylene blue on the TGT probe-modified Au-SPE during 25 times potential scans at the scan rate of 100 $mV.s^{-1}$ in a solution containing (v/v) 50% DMEM and 50% phosphate buffer (0.2 M, pH 7.4). (**b**) Cyclic voltammogram of methylene blue on the TGT probe-modified Au-SPE at various scan rates of 25, 50, 100, 150, 200, 250, and 300 $mV{\cdot}s^{-1}$ from inner to outer, in a solution containing (v/v) 50% DMEM and 50% phosphate buffer (0.2 M, pH 7.4). (**c**) A linear correlation was observed between both anodic (orange line) and cathodic (blue line) peak currents ($I_p$) and the changes in the scan rate (ʋ).

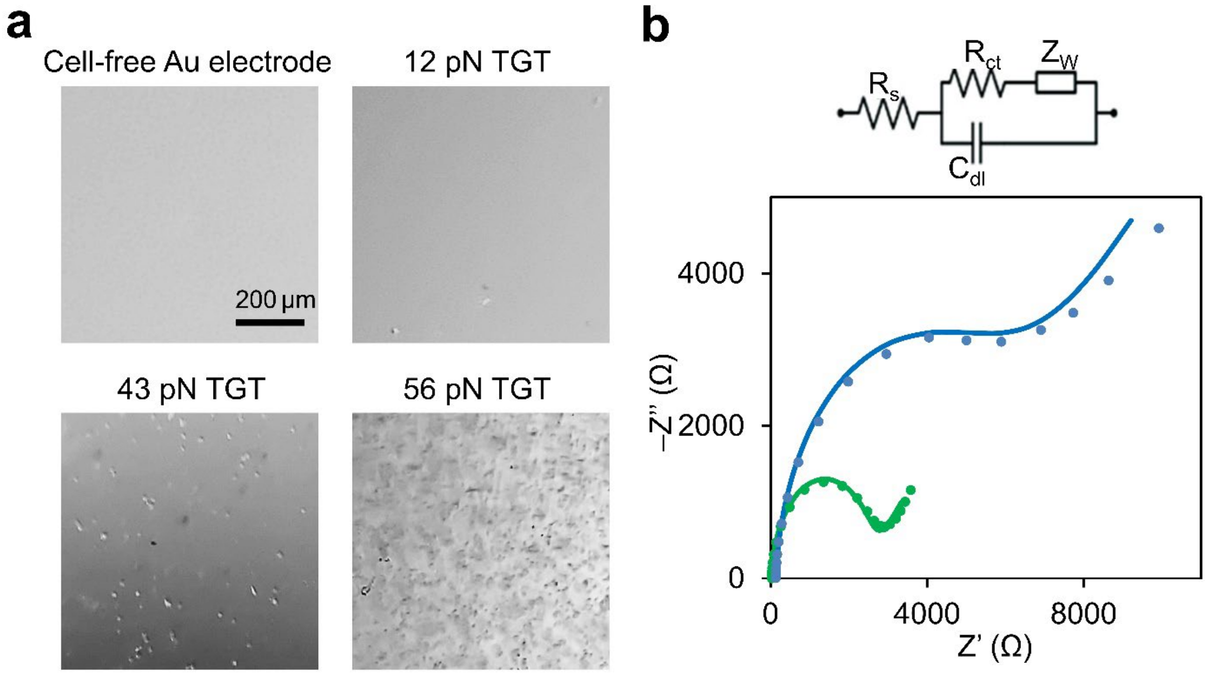


**Figure S4.** (**a**) Bright-field imaging of the TGT sensor before adding the HeLa cells or after a 60-min incubation with 100 μL of $1{\times}10^6$ HeLa cells/mL and washing respectively on 12 pN, 43 pN, and 56 pN TGT sensors. The sensors are made on the surface of a transparent gold electrode. (**b**) Nyquist plots of the 56 pN TGT sensor before (green line) and after (blue line) a 60-min incubation with 100 μL of $1{\times}10^6$ HeLa cells/mL and washing. The electrochemical impedance spectroscopy measurement was performed in a solution containing (v/v) 50% DMEM, 50% phosphate buffer (0.2 M, pH 7.4), and 5 mM ferrocene carboxylate, signals were recorded at an AC potential of 5 mV, a DC potential of 0.17 V, and in the frequency range of 100,000–0.1 Hz. The equivalent electric circuit compatible with the Nyquist diagrams were shown inset. $R_s$ is the solution resistance, $R_{ct}$ is the charge transfer resistance, $C_{dl}$ is double layer capacitance, and $Z_W$ is Warburg impedance.

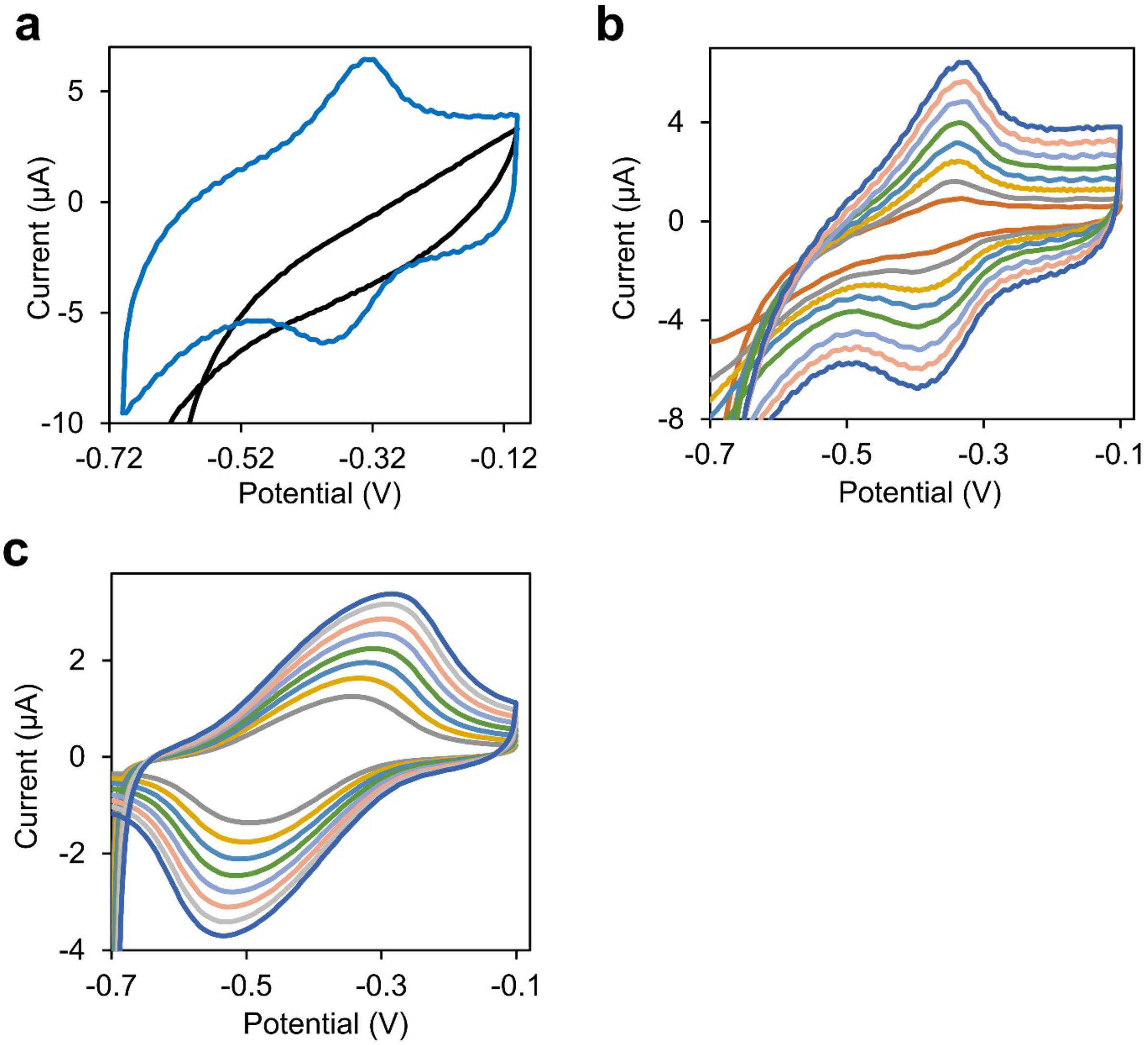


**Figure S5.** (**a**) Cyclic voltammogram of methylene blue on the surface of unmodified Au-SPE (black line) and the DNA hairpin force sensor-modified Au-SPE (blue line) as measured in a solution containing (v/v) 50% DMEM and 50% phosphate buffer (0.2 M, pH 7.4) at a scan rate of 400 mV·s$^{-1}$. (**b**) Cyclic voltammogram of methylene blue on the DNA hairpin force sensor-modified Au-SPE at various scan rates of 100, 150, 200, 250, 300, 350, 400, and 450 mV·s$^{-1}$ from inner to outer. (**c**) Cyclic voltammogram of methylene blue on the DNA hairpin force sensor-modified Au-SPE after adding 100 µL of 1×10$^{6}$ HeLa cells/mL for 60 min at various scan rates of 100, 150, 200, 250, 300, 350, 400, and 450 mV·s$^{-1}$ from inner to outer.

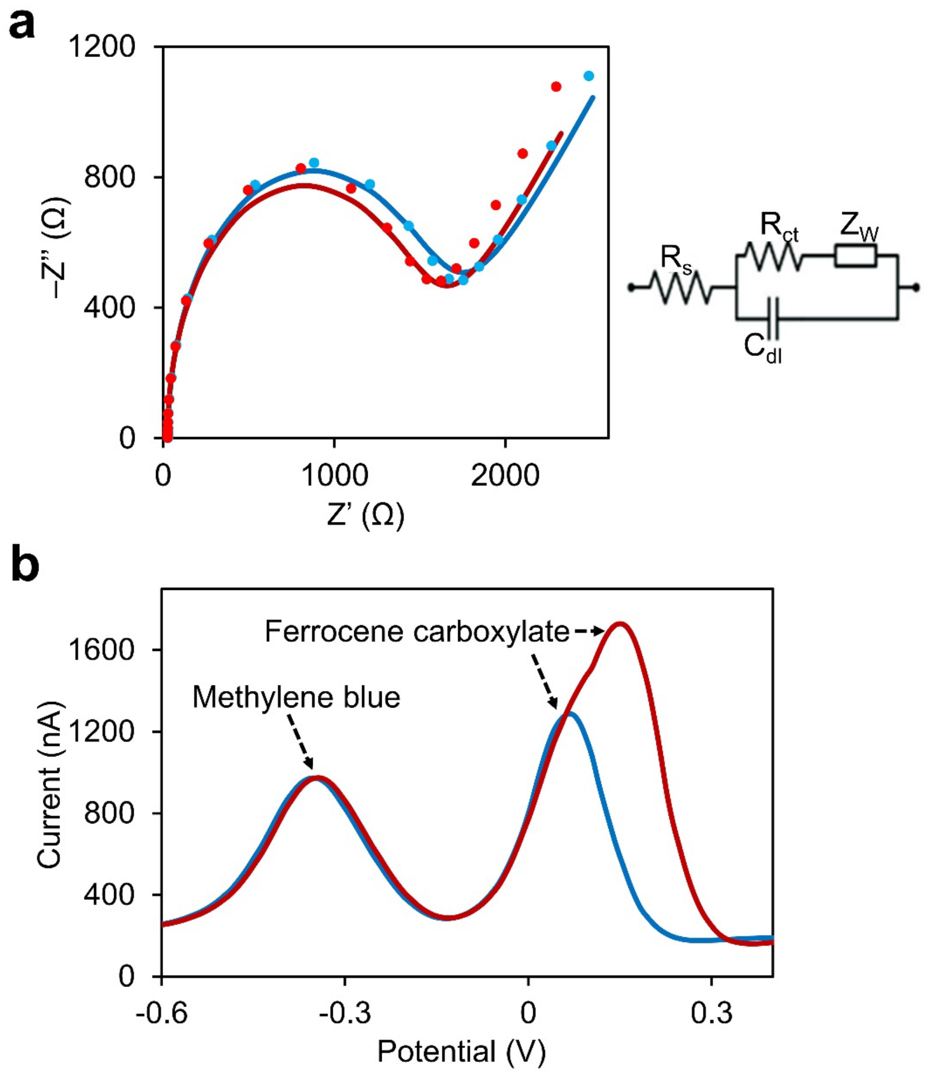


**Figure S6.** (**a**) Nyquist plots of a control Au-SPE that was immobilized with the DNA hairpin probe just without the attachment of biotinylated cRGDfK before (red line) and after (blue line) adding 100 μL of $1\times10^6$ HeLa cells/mL for 60 min. The electrochemical impedance spectroscopy measurement was performed in a solution containing (v/v) 50% DMEM, 50% phosphate buffer (0.2 M, pH 7.4), and 5 mM ferrocene carboxylate, signals were recorded at an AC potential of 5 mV, a DC potential of 0.17 V, and in the frequency range of 100,000–0.1 Hz. The equivalent electric circuit compatible with the Nyquist diagrams were shown on the right. $R_s$ is the solution resistance, $R_{ct}$ is the charge transfer resistance, $C_{dl}$ is double layer capacitance, and $Z_W$ is Warburg impedance. (**b**) Square wave voltammetry of another control Au-SPE before (red line) and after (blue line) adding 100 μL of $1\times10^4$ HeLa cells/mL for 60 min. This control electrode was directly immobilized with cRGDfK ligands and cRGDfK-free DNA hairpin probe. The measurement was performed in a solution containing (v/v) 50% DMEM, 50% phosphate buffer (0.2 M, pH 7.4), and 10 μM ferrocene carboxylate. The step potential was set as 20 mV, the pulse amplitude was at 50 mV, and the frequency was at 20 Hz.
